# GLP-1 receptor agonist ameliorates obesity-induced chronic kidney injury via restoring renal lipid and energy metabolism homeostasis: revealed by metabolomics

**DOI:** 10.1101/194654

**Authors:** Chengshi Wang, Ling Li, Shuyun Liu, Guangneng Liao, Younan Chen, Jingqiu Cheng, Yanrong Lu, Jingping Liu

**Author notes:** Corresponding Authors: Jingping Liu, Yanrong Lu, Address: Key Laboratory of Transplant Engineering and Immunology, West China Hospital, Sichuan University, No. 1 Keyuan 4^th^ Road, Gaopeng Ave, Chengdu 610041, China.

## Abstract

Increasing evidence indicate that obesity is highly associated with chronic kidney disease (CKD).GLP-1 receptor (GLP-1R) agonist has shown benefits on kidney diseases, but its direct role on kidney metabolism in obesity is still not clear. This study aims to investigate the protection and metabolic modulation role of liraglutide (Lira) on kidney of obesity. Rats were induced obese by high-fat diet (HFD), and renal function and metabolism changes were evaluated by metabolomic, biological and histological methods. HFD rats exhibited metabolic disorders including elevated body weight, hyperlipidemia and impaired glucose tolerance, and remarkable renal injuries including declined renal function and inflammatory/fibrotic changes, whereas Lira significantly ameliorated these adverse effects in HFD rats. Metabolomic data showed that Lira reduced renal lipids including fatty acid residues, cholesterol, phospholipids and triglycerides, and improved mitochondria metabolites such as succinate, citrate, taurine, fumarate and NAD^+^ in the kidney of HDF rats. Furthermore, we revealed that Lira inhibited renal lipid accumulation by coordinating lipogenic and lipolytic signals, and rescued renal mitochondria function via Sirt1/AMPK/PGC1α pathways in HDF rats. This study suggested that Lira alleviated HFD-induced kidney injury via directly restoring renal lipid and energy metabolism, and GLP-1 receptor agonist is a promising therapy for obesity-associated CKD.

## Introduction

Chronic kidney disease (CKD), characterized by progressive destruction of renal mass and slow loss of renal function, is the leading cause to end-stage kidney disease (ESKD) and renal failure (Bolignano and Zoccali, 2013). Obesity is an epidemic disease and contributes to type 2 diabetes and cardiovascular diseases with high mortality (Goran et al., 2003). However, increasing data indicate that obesity is also highly linked with the onset and progression of CKD (Fox et al., 2004; Gelber et al., 2005; Wang et al., 2008). Epidemiologic studies reported that obesity was associated with a 70% increased risk of microalbuminuria compared to lean subjects (Stenvinkel et al., 2013). High body mass index (BMI) has been considered as one of the strongest risk factors for new-onset CKD (Fox et al., 2004), and higher baseline BMI remained an independent predictor for ESRD after adjustments for BP and DM (Hsu et al., 2006). Obesity may initiate the slow development of kidney dysfunction and lesions including mesangial expansion, glomerular hypertrophy and glomerulosclerosis (Mathew et al., 2011). The mechanism of obesity-induced kidney injury is complex and multiple factors such as oxidation stress, inflammation, TGF-β, lipotoxicity, renin-angiotensin system and adipocyte-derived hormones are involved (Mathew et al., 2011; Thethi et al., 2012). As the increased prevalence of obesity over the world, it is of great clinical benefits to find therapies for obesity-associated CKD.

Glucagon-like peptide-1 (GLP-1) is a polypeptide hormone secreted from intestines, which can regulate blood glucose level in diabetic patients, and promote pancreatic cells proliferation in DM models (Buteau, 2008; Doyle and Egan, 2007). GLP-1 exerts its biological actions via binding to its receptor (GLP-1R) on the surface of various cells ^11^. Besides to pancreatic β-cells, GLP-1R has also been found in kidney cells such as glomerular endothelium and mesangial cells (Mima et al., 2012; Takashima et al., 2016). Previous studies have proved the benefits of GLP-1R agonist therapy on acute and chronic kidney diseases (Katagiri et al., 2013; Mima et al., 2012). GLP-1R agonist exendin-4 (Ex-4) ameliorated cisplatin-induced acute renal tubular cell injury and apoptosis, while inhibition of GLP-1R abolished its renoprotective effect (Katagiri et al., 2013). GLP-1R agonist liraglutide (Lira) alleviated diabetic nephropathy (DN) in mice through inhibiting glomerular superoxide and NAD(P)H oxidase (Fujita et al., 2014). Recent study further reported that GLP-1R agonist Ex-4 inhibited renal cholesterol accumulation and inflammation in diabetic apoE knockout mice (Yin et al., 2016). However, the direct role of GLP-1R agonist on renal metabolic abnormalities in obesity remains not clear.

In this study, we aim to evaluate the renal protection effect of GLP-1R agonist (Lira) in the obese rats induced by HFD. By using metabolomic, biological and histological methods, we further revealed the metabolic modulation role of GLP-1 on obesity-induced kidney injury.

## Materials and methods

### Ethical Statement

All animal work were conducted under the approved guidelines of Sichuan University (Chengdu,China) and approved by the Animal Care and Treatment Committee of Sichuan University (Chengdu, China). The methods were performed in accordance with approved guidelines.

### Animal experiment

Male Sprague-Dawley (SD) rats were purchased from Experimental Animal Center of Sichuan University (Chengdu, China). Animals were housed in cages with controlled temperature, humidity and 12 h cycles of light and darkness, and fed with standard chow and tap water ad libitum. Rats were randomly divided into control (n = 10), HFD (n = 10) and HFD + Lira (n = 10) group. Control rats were fed with regular diet, and HFD rats were fed with high-fat diet (regular diet plus 20% lard, 15% sucrose, 1.5% cholesterol, 0.1% sodium cholate). At 12 weeks after HFD, rats in HFD + Lira group were twice-daily subcutaneously injected with Lira (0.1 mg/kg, Novo Nordisk, Denmark) for 12 further weeks, and rats in HFD group were given PBS. The samples of blood, urine and kidney from rats were collected at 24 weeks after HFD experiment beginning.

### Biochemical measurement

Clinical biochemistry analysis was performed on an Automatic Biochemistry Analyzer (Cobas Integra 400 plus, Roche) by commercial kits with the following parameters: blood glucose (BG), cholesterol (TC), triglyceride (TG), low-density lipoprotein cholesterol (LDL-C), high-density lipoprotein cholesterol (HDL-C), blood urea nitrogen (BUN), creatinine (CREA) and urinary albumin to creatinine ratio (UACR).

### Oil Red O (ORO) staining

Fresh renal tissues obtained from rats were made to 5 μm frozen sections, and then incubated with Oil Red O solution (KeyGen Biotech Co. Ltd., Nanjing, China) at room temperature for 30 min. After washing twice with PBS, the stained sections were observed on a light microscope (Carl Zeiss, Germany).

### Measurement of lipid peroxidation

Renal lipid peroxidation was measured by malondialdehyde (MDA, Beyotime Institute of Biotechnology, China) kit according to the manufacturer’s instructions. Briefly, the supernatant of renal homogenate was added to TBA solution and incubated at 100 °C for 15 min, and then placed on ice to terminate the reaction. After centrifugation, the supernatant of reaction solution was measured by microplate reader (BioTek, MQX200, USA) at 535 nm.

### ^1^H NMR analysis of kidney tissue

Renal tissues were extracted by chloroform/methanol method as previously described ^34^. The aqueous extracts were lyophilized and dissolved in 500 μl PBS containing 50% (v/v) D_2_O and 0.001% (w/v) 3-trimethylsilyl propionic acid-d4 sodium salt (TSP, Cambridge Isotope Laboratories, USA). The lyophilized lipid extracts were dissolved in 500 μl deuterated chloroform (CDCl_3_) containing 0.03% (w/v) tetramethylsilane (TMS). NMR analysis was performed on a Bruker Avance II 600-MHz spectrometer (Bruker Biospin, Germany) at 298 K with a 5-mm PATXI probe. The ^1^H NMR spectra of renal extracts was acquired by a NOESY-presaturation (NOESYGPPR1D) pulse sequence (RD-90°-t1-90°-tm-90°-acq) with following parameters: relaxation delay (RD) = 5 s, mixing time = 100 ms, acquisition time = 2.48 s. The 90° pulse length was about 10 μs, and 64 scans were collected into 32 K data points with a spectral width of 11 ppm. The FIDs were weighted by an exponential function with a 0.3 Hz line broadening and zero-filled to 32 k data points prior to fourier transform (FT). The chemical shifts were referenced to TSP or TMS at 0 ppm.

### Multivariate data analysis

NMR spectra were manually phased and baseline corrected, and then binned into equal widths (0.003 ppm) corresponding to the 0.5-9.5 ppm (aqueous extract) and 0.5-5.5 ppm (lipid extract) regions by MestReC software (version 4.9.9.9, Mestrelab Research, Spain). The integrals were normalized to the sum intensity of each spectrum excluding the water (4.7-5.2 ppm) regions. The normalized bins were then subjected to principal component analysis (PCA) and orthogonal projection to latent structure with discriminant analysis (OPLS-DA) by SIMCA-P (version 11.5, Umetrics, Sweden). OPLS-DA models were validated by a seven-fold cross validation method with a permutation test to avoid over-fitting. The OPLS-DA coefficient plot was generated with an in-house developed MATLAB script as previously described (Liu et al., 2015).

### Mitochondrial ROS (mtROS) assay

The kidney tissue mtROS was measured by mitochondrial superoxide indicator (MitoSOX, Molecular Probes, Thermo Fisher Scientific, USA) according to the manufacturer’s instructions. Briefly, fresh renal tissues were made to 5 μm frozen sections, and then incubated with MitoSOX solution (2 μM) at 37°C for 10 min. After washing with PBS, the stained sections were observed on a fluorescent microscope (Axioplan 2, Carl Zeiss, Germany).

### Measurement of kidney ATP

Renal tissue ATP was measured by Luciferase ATP Assay Kit (Beyotime, China) according to the manufacturer’s instructions. Briefly, 20 mg tissues were lysed with 200 μl lysis buffer on ice bath, and then centrifuged 12000 g for 5 minutes at 4 °C. 100 μl of supernatant was collected and mixed with 100 μl ATP detection solution. Luminance was measured by fluorescence microplate reader (Synergy4, BioTek, USA). The ATP level was normalized to the protein concentration of each sample.

### Quantitative real-time PCR (qPCR)

Total RNA was isolated from renal tissues by using Trizol (GIBCO, Life Technologies, USA) according to the manufacturer’s instructions. Total RNA was quantified by NanoDrop 2000 spectrophotometer (Thermo Fisher Scientific Inc., USA). cDNA was synthesized by an iScript cDNA Synthesis Kit (Bio-Rad, CA, USA). The sequences of primers were listed in Table S1. qPCR analysis were performed on a CFX96 Real-Time PCR Detection System (Bio-Rad, CA, USA) with SYBR green supermix (SsoFast EvaGreen, Bio-Rad, USA). The change folders of mRNAs were calculated by delta-delta Ct method with β-actin as internal reference gene.

### Histological examination

The fixed renal tissues were embedded in paraffin and made to 5 μm sections. Renal sections were deparaffinized in xylene and rehydrated in graded ethanols, and then stained with hematoxylin eosin (HE) and periodic acid-schiff (PAS) staining. For immunohistochemical (IHC) staining, sections were blocked with 1% BSA, and incubated with diluted primary antibodies including rabbit anti-α-SMA (Abcam, USA), mouse anti-MCP-1 (Abcam, USA), rabbit anti-IL-6 (Abcam, USA), rabbit anti-CPT1 (Proteintech, USA), rabbit anti-FABP1 (Proteintech, USA), rabbit anti-PPARα (Proteintech, USA), rabbit anti-UCP2 (Abcam, USA), rabbit anti-ATP5A1 (ABClonal, USA), rabbit anti-Sirt1 (CST, USA), rabbit anti-p-AMPK (CST, USA) and rabbit anti-PGC1α (Proteintech, USA), then incubated with HRP-conjugated secondary antibody (DAKO, USA), and finally stained with DAB substrate and hematoxylin. The images of stained sections were acquired by microscope (Carl Zeiss, Germany), and quantitative analysis of positive staining areas (%) in images was done by using Image J software (NIH, USA).

### Statistical analysis

Descriptive statistics were presented as mean ± SD and analyzed by SPSS software (version 11.5, IBM Corp., USA). Comparison among groups was analyzed with one-way analysis of variance (ANOVA), and p < 0.05 was considered as significant difference.

## Results

### The effect of Lira on general and clinical parameters in HFD rats

As shown in Table 1, HDF rats had higher levels of body weight (BW), BG, TC, TG, LDL-C and areas under the curve (AUC) of IPGTT (Figure S1), and lower level of HDL-C compared to control. By contrast, Lira significantly reduced BW, BG, AUC and blood lipid levels in HFD rats. In addition, HFD rats showed increased levels of BUN, CREA and UACR compared to control, while Lira significantly reduced CREA and UACR in HFD rats.

**Table 1.** General and biochemical parameters of rats in different groups

| Parameters | Control | HDF | HDF + Lira |
| --- | --- | --- | --- |
| BW (g) | 451.8±37.32 | 625.8±45.72** | 512.6±49.31<sup>#</sup> |
| BG (mmol/L) | 5.60±1.59 | 9.98±3.46** | 7.45±2.69<sup>#</sup> |
| TC (mmol/L) | 0.70±0.20 | 1.82±0.34** | 1.12±0.14<sup>##</sup> |
| TG (mmol/L) | 0.26±0.11 | 0.51±0.27** | 0.33±0.14<sup>#</sup> |
| LDL-C (mmol/L) | 0.18±0.06 | 0.98±0.33** | 0.67±0.22<sup>#</sup> |
| HDL-C (mmol/L) | 1.03±0.13 | 0.55±0.19** | 0.79±0.24<sup>#</sup> |
| BUN (mmol/L) | 4.24±0.86 | 5.88±1.92* | 4.97±1.03 |
| CREA (μmol/L) | 35.18±10.95 | 76.66±21.43** | 54.68±17.32<sup>#</sup> |
| UACR (mg/mmol) | 2.34±0.73 | 8.80±3.15** | 5.38±1.69<sup>#</sup> |

Note: \* p<0.05, \*\* p<0.01 compared with control; # p<0.05, <sup>##</sup> p<0.01 compared with HFD.

### Lira reduced renal inflammation and fibrosis in HFD rats

HFD rats showed obvious renal lesions including tubular hypertrophy, basement membrane thickening, mesangial expansion and glomerulosclerosis compared to control rats (Figure 1A). In contrast, Lira markedly reduced kidney histological injury and fibrosis in HFD rats. IHC staining results showed that HFD rats had increased levels of MCP-1, IL-6 and α-SMA, while Lira inhibited these pro-inflammatory and pro-fibrotic factors expression in the kidney of HFD rats (Figure 1B-C).

**Figure 1.**
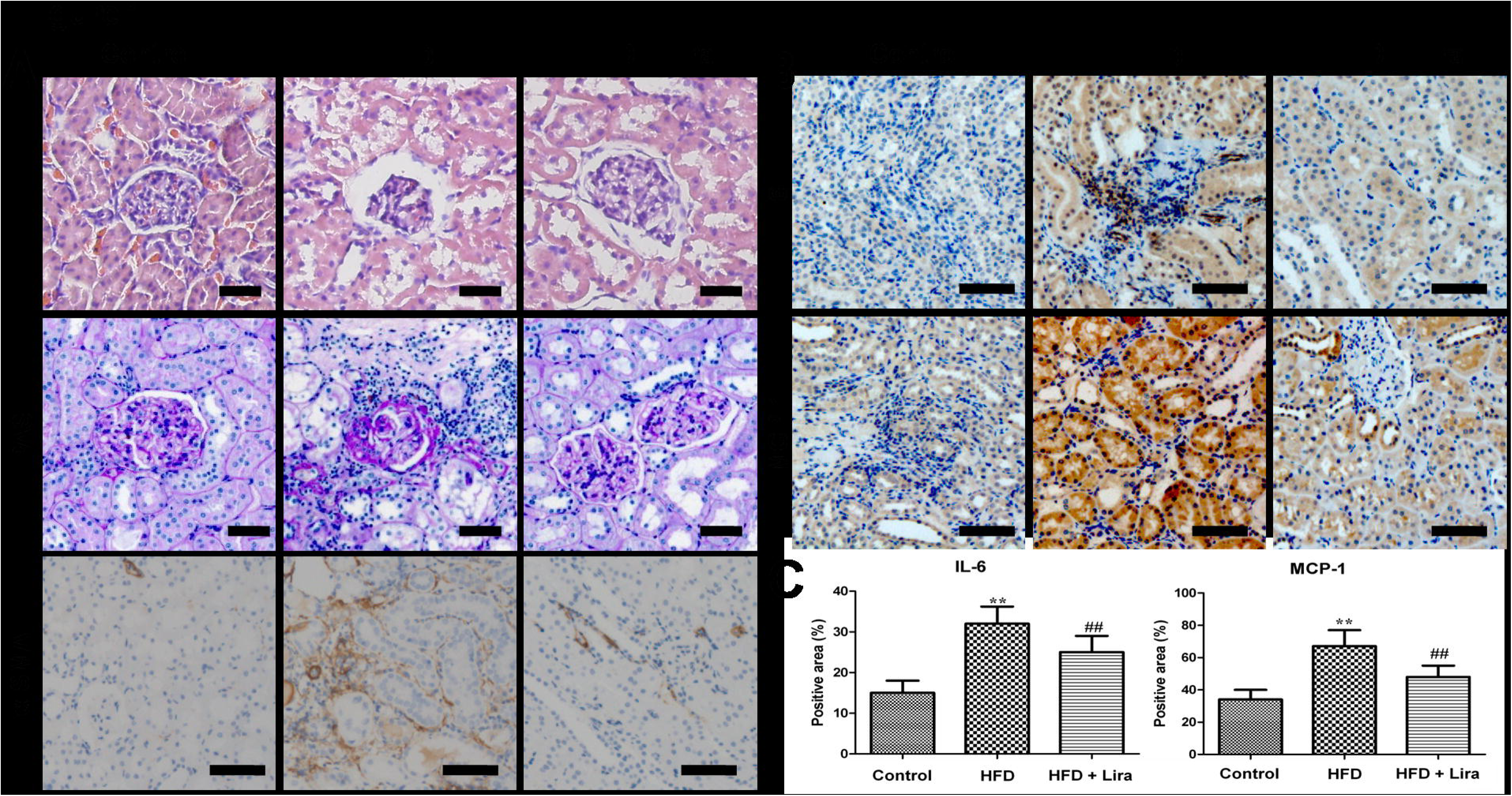
(A) Representative HE, PAS staining (scale bar = 50 μm) and IHC staining for α-SMA (scale bar = 100 μm) of kidney from control, HFD and HFD + Lira group. (B) IHC staining for IL-6 and MCP-1 in kidney of different groups (scale bar = 100 μm). (C) Quantification of IL-6 and MCP-1 protein detected by IHC staining (** p<0.01, compared with control; ^#^ p<0.05, ^##^ p<0.01, compared with HFD).

### Lira ameliorated renal metabolic disorders in HDF rats

The representative ^1^H NMR spectra of lipid and aqueous extract from rat kidney were shown in Figure S2 and S3, respectively. OPLS-DA score plots showed clear clustering among the three groups (Figure 2 and 3). In lipid extract, HFD rats showed higher level of fatty acid residues, cholesterol, phospholipids and triglycerides compared to controls, while Lira reduced these lipids levels in the kidney of HFD rats (Figure 2). In aqueous extract, HFD rats showed lower level of leucine/isoleucine, valine, lysine, glutamine, glutamate, succinate, citrate, taurine, glycine, tyrosine, histidine, phenylalanine and NAD^+^, and higher level of alanine, creatinine, choline, allantoin and formate compared to controls. In contrast, Lira reduced the level of alanine, choline, glucose and allantoin, and increased the level of lysine, acetate, succinate, citrate, taurine, fumarate, histidine and NAD^+^ in the kidney of HFD rats (Figure 3).

**Figure 2.**
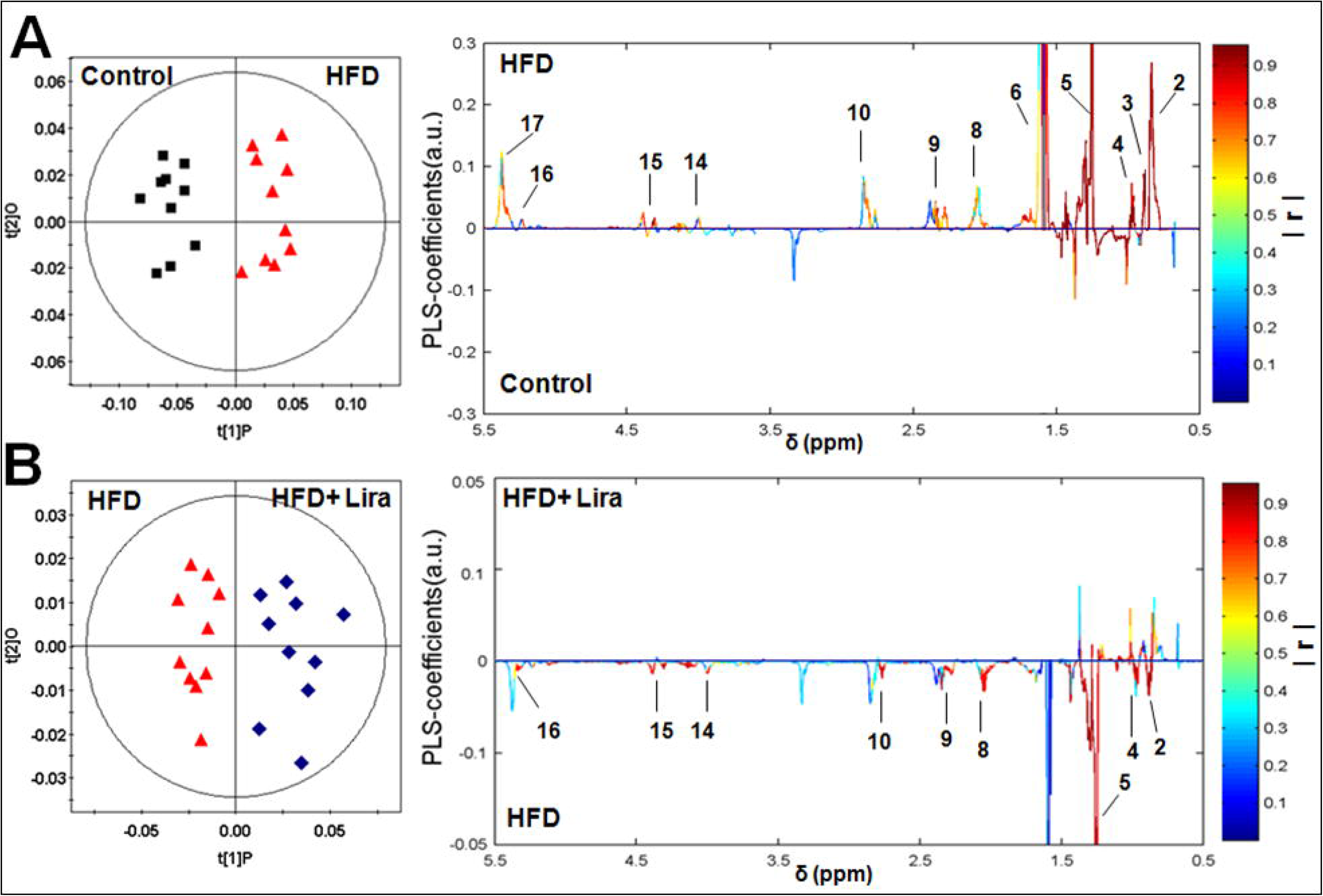
OPLS-DA loading plots derived from renal lipid extracts of ^1^H NMR spectra in (A) HFD group *vs* control group and (B) HFD + Lira group *vs* HFD group. Note: 1. Total Cholesterol (C_18_H_3_), 2. Fatty acid residues (ω-CH_3_), 3. Total Cholesterol (C_26_H_3_, C_27_H_3_), 4. Free Cholesterol (C_19_H_3_), 5. Fatty acid residues ((CH_2_-)_n_), 6. Fatty acid residues (COCH_2_-CH_2_), 7. Fatty acid residues (-CH_2_ of ARA+EPA), 8. Fatty acid residues (CH_2_-CH=), 9. Fatty acid residues (-CO-CH_2_), 10. Fatty acid residues (-CH=CH-CH_2_-CH=CH-), 11. Phosphatidylethanolamine (-CH_2_-NH_2_), 12. Sphingomyelin (-CH_2_-N-(CH_3_)_3_), 13, Phosphatidylcholine (-CH_2_-N-(CH_3_)_3_), 14. Total phospholipids (Glycerol (C_3_H_2_)), 15. Triglycerides (C_1_H and C_3_H of glycerol), 16. Triglycerides (C_2_H of glycerol), 17. Fatty acid residues (-CH=CH-).

**Figure 3.**
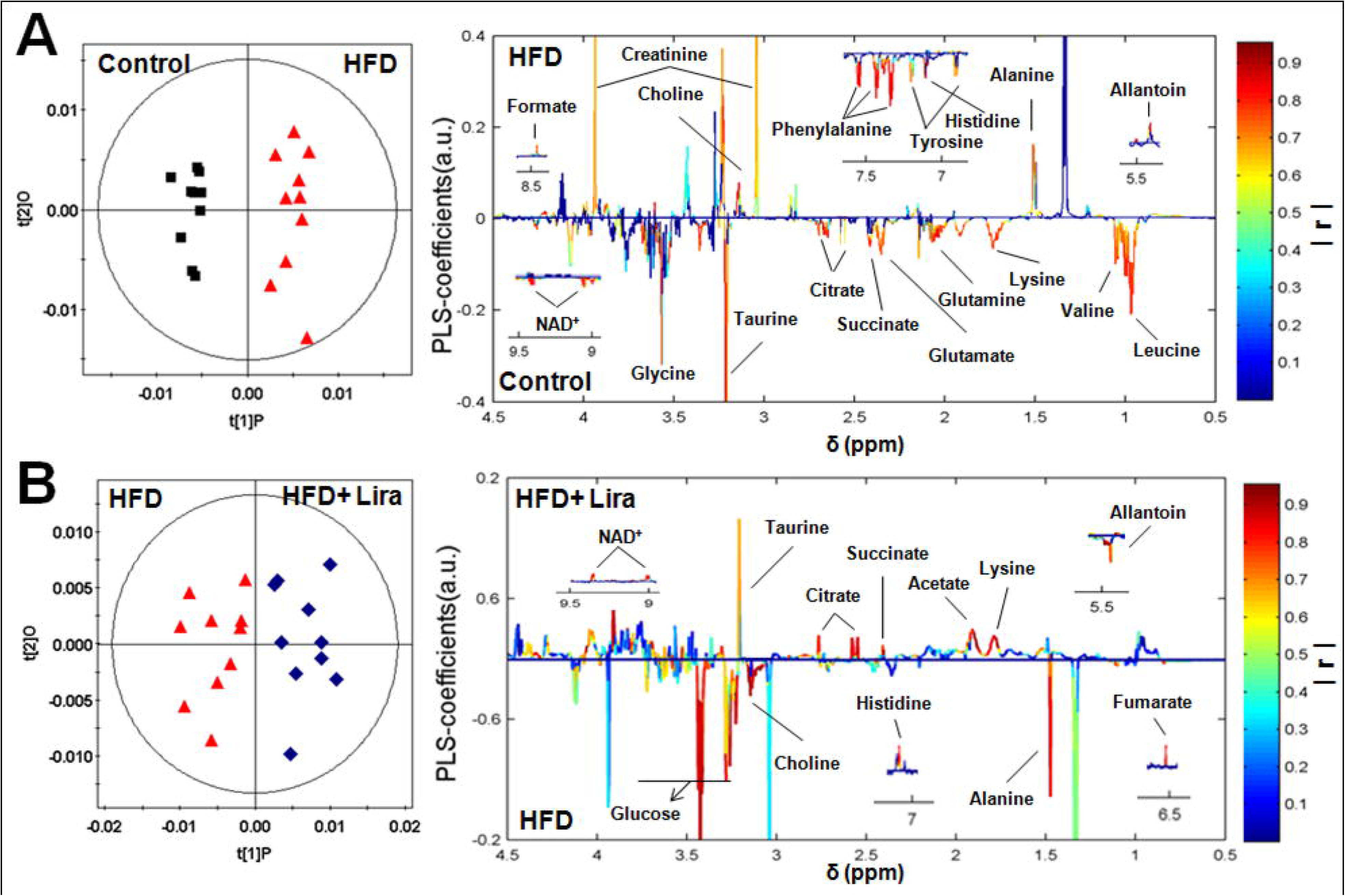
OPLS-DA loading plots derived from renal aqueous extracts of ^1^H NMR spectra in (A) HFD group *vs* control group and (B) HFD + Lira group *vs* HFD group. The colored correlation coefficients indicated significant metabolites which were higher or lower in each group.

### Lira decreased renal lipids accumulation in HDF rats

Compared with controls, HFD rats showed increased lipid droplets formation (Figure 4A) and MDA (Figure 4B) in the kidney, while Lira reduced the lipid accumulation and lipid peroxidation in the obese kidney. In addition, HFD rats showed increased level of lipogenic genes including CD36, L-FABP, SREBP-1c and FAS, and reduced lipolytic genes including CPT-1 and PPARα. In contrast, Lira significantly reduced these lipogenic genes and enhanced lipolytic genes expression in the kidney of HFD rats (Figure 4C). IHC staining results showed that Lira reduced FABP1 and increased PPAR-α and CPT1 protein level in kidney of HFD rats (Figure 4D-E).

**Figure 4.**
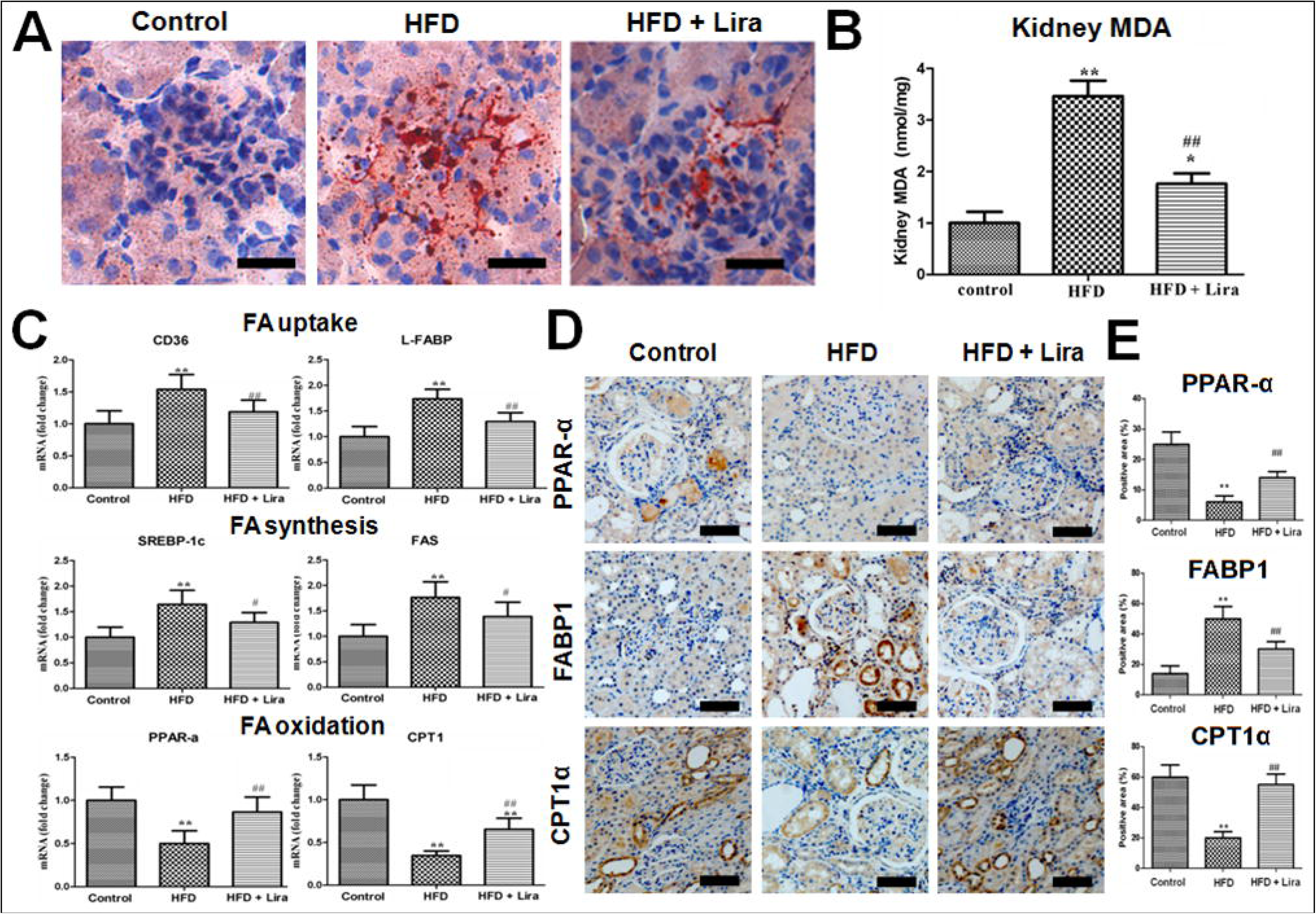
(A) Oil Red O (ORO) staining of kidney from control, HFD and HFD + Lira group (scale bar = 100 μm). (B) Quantification of kidney MDA level. (C) Real-time PCR analysis for CD36, L-FABP, SREBP-1c, FAS, PPAR-α and CPT1 mRNA. (D) IHC staining for renal PPAR-α, FABP1 and CPT1 (scale bar = 100 μm). (E) Quantification of PPAR-α, FABP1 and CPT1 protein detected by IHC staining (* p<0.05, ** p<0.01, compared with control; ^#^ p<0.05, ^##^ p<0.01, compared with HFD).

### Lira restored renal mitochondrial function in HDF rats

As shown in Figure 5, HFD rats showed increased renal level of mtROS (Figure 5A-B) and reduced level of ATP (Figure 5C), while Lira significantly inhibited mtROS and promoted ATP production in the kidney of HFD rats. In addition, Lira reversed the decline of mitochondria biogenesis genes including TFAM, NRF-1, NDUFS5 and SDHB expression in kidney of HFD rats (Figure 5D). IHC staining results showed that Lira increased ATP5a1 and UCP2 protein levels in the kidney of HFD rats (Figure 5E-F)

**Figure 5.**
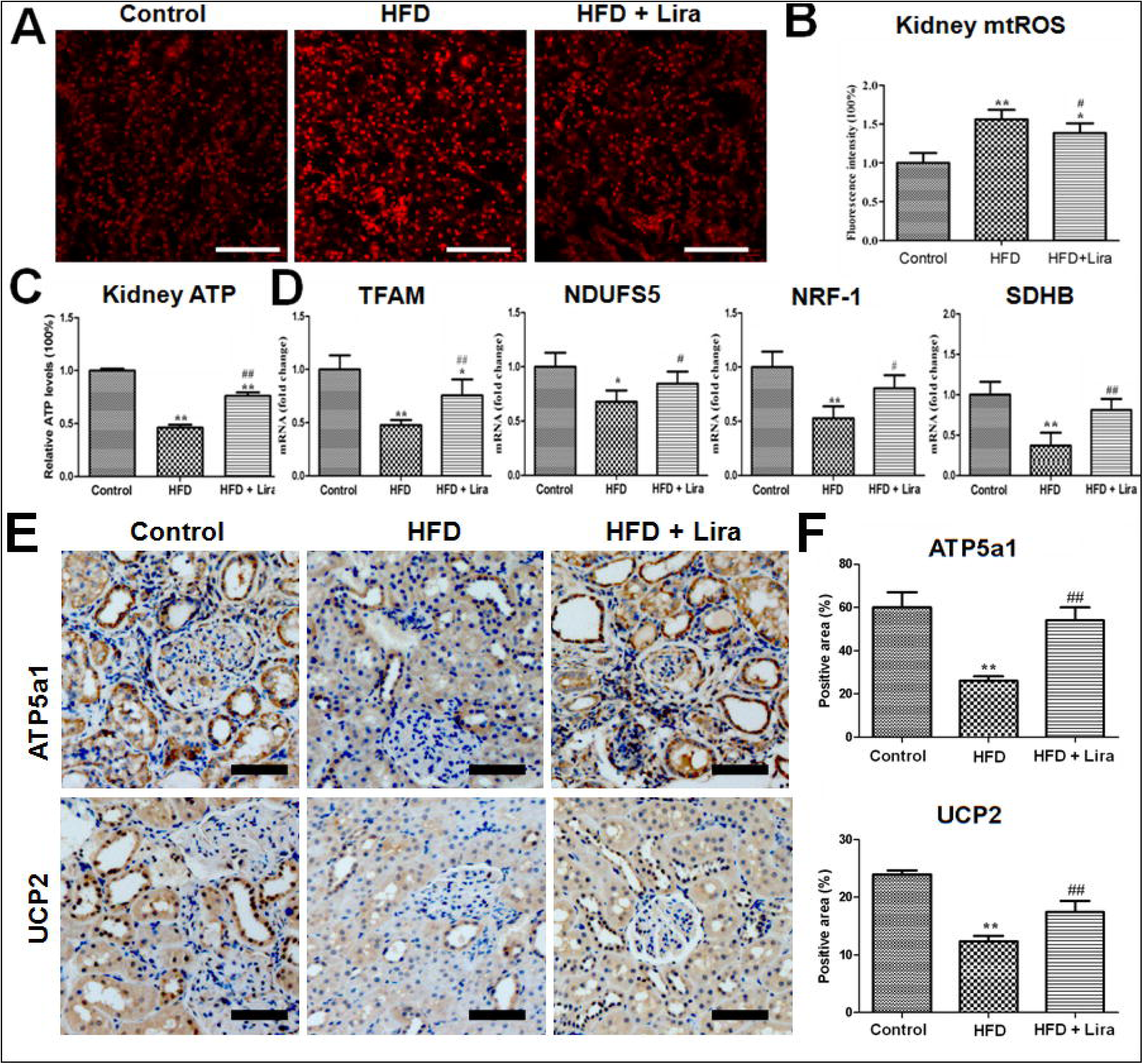
(A) Fluorescence staining of MitoSOX in kidney from control, HFD and HFD + Lira group (scale bar = 100μm). (B) Quantification of kidney mtROS detected by MitoSOX staining.(C) Renal tissue ATP levels in different groups. (D) Real-time PCR analysis for TFAM, NRF-1, NDUFS5 and SDHB mRNA. (E) IHC staining for renal UCP2 and ATP5a1 (scale bar = 100μm).(F) Quantification of UCP2 and ATP5a1 protein detected by IHC staining (* p<0.05, ** p<0.01 compared with control; ^#^ p<0.05, ^##^ p<0.01 compared with HFD).

### Lira activated renal Sirt1/AMPK/PGC1α signals in HDF rats

As shown in Figure 6B, qPCR results showed that HFD rats had reduced mRNA levels of Sirt1 and PGC1α in kidney, while Lira significantly up-regulated these genes expression in the kidney of HFD rats. In addition, IHC staining results showed that HFD inhibited Sirt1, PGC1α and p-AMPK expression in kidney, whereas Lira significantly increased these proteins expression in the kidney of HFD rats (Figure 6A and C).

**Figure 6.**
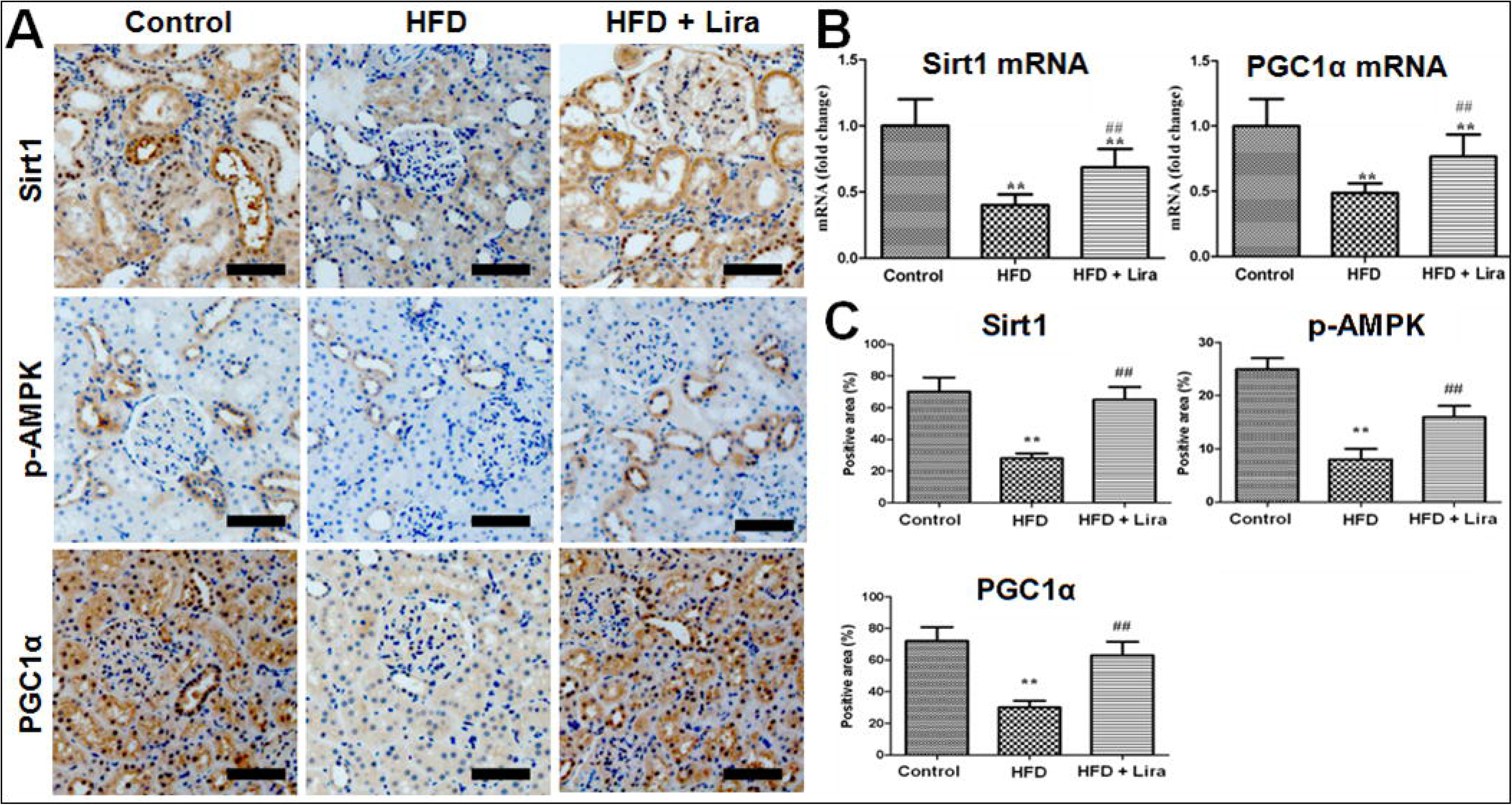
(A) IHC staining for PGC1α, p-AMPK and Sirt1 of kidney from control, HFD and HFD+ Lira group (scale bar = 100μm). (B) Real-time PCR analysis for Sirt1 and PGC1α mRNA. (C) Quantification of PGC1α, p-AMPK and Sirt1 protein detected by IHC staining (** p<0.01 compared with control; ^##^ p<0.01 compared with HFD).

## Discussion

It is well-documented that high-fat diets (HFD) increased the prevalence of obesity and its associated metabolic syndrome, T2DM and cardiovascular diseases ^2^. GLP-1R agonist had been proved to reduce food intake, body weight and blood glucose in obese rats (Raun et al., 2007). Consisted with previous report, we observed that Lira reversed the general metabolic disorders including elevated body weight, hyperlipidemia and impaired glucose tolerance induced by HFD.

Moreover, HFD rats showed remarkable kidney lesions including increased levels of renal function parameters, oxidative stress, inflammatory and fibrotic changes, and these factors have been well recognized as key mediators for the development of CKD. The association between obesity and CKD has been already purposed, and HFD increased oxidative stress markers such as 8-OH-dG, and induced sustained inflammation and cell apoptosis in the kidney (Chung et al., 2012). Previous study found that GLP-1R expressed on the renal glomeruli and endothelial cells, and GLP-1R therapy had beneficial effects on diabetic kidney diseases (Einbinder et al., 2016).

GLP-1R agonist Ex-4 could attenuate renal oxidative stress, ECM deposition and cytokines release in STZ-induced DN rats via direct effects on the GLP-1R in kidney tissue (Kodera et al., 2011). In this study, we observed that Lira significantly improved renal function, and ameliorated renal histological injury, pro-inflammatory and pro-fibrotic factors expression in HFD rats. These results again demonstrated that GLP-1R agonist Lira had renal protective role in obese rats.

The anti-inflammatory and antioxidant effects of GLP-1R agonist on kidney injury are well-known (Einbinder et al., 2016; Kodera et al., 2011; Mima et al., 2012), but its direct effect on renal metabolism in obesity has rarely reported. To further explore the metabolic mechanism of Lira for kidney protection, the renal metabolic profile of HFD rats was measured by ^1^H NMR-based metabolomics. Our results showed that HFD rats had higher levels of renal lipids including fatty acid residues, cholesterol, phospholipids and triglycerides, while Lira prevented these lipids deposition in the kidney of HFD rats. Furthermore, Lira reversed the decline of succinate, citrate, taurine, fumarate and NAD^+^ in the kidney of HFD rats. It is well-documented that succinate, citrate and fumarate were converted in TCA cycle of mitochondria (Sharma et al., 2013), and thus their reduction reflected the disturbed energy metabolism and mitochondria dysfunction in kidney of HFD rats. In addition, NAD^+^ is an important substrate required for oxidative phosphorylation and mitochondrial respiration, and reduced NAD^+^ has been linked to oxidative stress, energy imbalance and down-regulation of sirtuins in obesity, metabolic syndrome and diabetes (Imai and Guarente, 2014). Therefore, our results suggested that lipid and energy metabolism were involved in the renal protection of Lira on HFD rats.

Previous studies had found abnormal lipid deposition in the kidney of metabolic diseases such as obesity and diabetes, and suggested that lipotoxicity contributes to the development of kidney injury such as glomerulopathy, tubulointerstitial inflammation, tubular cell apoptosis and renal fibrosis (Weinberg, 2006). Fatty acids (FAs) are key mediators of lipotoxicity, and their levels are dramatically elevated in obese patients (Bobulescu, 2010). The abnormally accumulated FAs lead to kidney injury through complex mechanism including ROS, ER stress, mitochondria dysfunction and activation of pro-inflammatory and pro-fibrotic pathways (Nosadini and Tonolo, 2011). In this study, we found that Lira significantly reduced the renal lipids including FAs accumulation in HFD rats, suggesting that Lira might directly regulate FA metabolism in obese kidney. SREBP-1 is a master regulators of lipid metabolism, which promotes lipogenesis by inducing target genes involved in the FA transport (CD36, FABP1) and FA biosynthesis (ACC and SCD-1) (Frederico et al., 2011; Nosadini and Tonolo, 2011). In contrast, PPARα improves lipolysis via increasing genes related to FA oxidation such as CPT-1 (Wang et al., 2014). Our results further showed that Lira inhibited SREBP-1c, FAS, CD36 and FABP1, and increased CPT-1 and PPARα level in the kidney of HDF rats. Thus, these results suggested that Lira alleviated kidney injury in obese rats at least partly by directly reducing renal lipid dysfunction.

The kidney is a high energy demand organ that can use various substrates as fuels to generate ATP. In renal tubule cells, mitochondrial oxidation of FFA is a major source for ATP production due to their little glycolytic capacity (Bobulescu, 2010). Mitochondria play a centre role in mediating cellular energy metabolism, while obesity exhibited energy depletion and mitochondrial dysfunction associated with declined key mitochondria regulator proteins including PGC1α and AMPK (Cheng and Almeida, 2014; Fox et al., 2004; Jelenik and Roden, 2013). Previous reports indicated that renal accumulated lipids caused mitochondria stress including increased mtROS, depolarized mitochondrial membrane potential and impaired ATP generation in kidney cells (Decleves et al., 2014; Xu et al., 2015). In this study, we observed that Lira prevented renal mitochondria injury induced by HFD, as demonstrated by reduced mtROS and increased level of TCA metabolites, ATP and mitochondria biogenesis genes including TFAM, NRF-1, NDUFS5 and SDHB, and increased ATP5a and UCP2 protein in kidney of HFD rats. Sirt1 is a NAD^+^ dependent deacetylase, which plays a key role in regulating mitochondria oxidative metabolism and energy homeostasis (Imai and Guarente, 2014). Sirt1 had been reported to regulate energy metabolism and mitochondria biogenesis through a complex network involving AMPK and PGC1α (Canto and Auwerx, 2009). It had been reported that GLP-1R agonist exenatide activated Sirt1/AMPK pathways and thus reduced lipid deposition and inflammatory response in the liver of HFD mice(Xu et al., 2014). We also observed decline of renal NAD^+^ and Sirt1 in HFD rats, whereas Lira increased level of NAD^+^ and Sirt1/p-AMPK/PGC1α expression in the kidney of HFD rats. There results suggested that Lira prevented kidney injury by restoring mitochondrial function and activating Sirt1/AMPK/PGC1α pathways in HFD rats.

## Conclusion

In summary, HFD-induced obese rats showed impaired renal function and elevated renal inflammation, oxidative stress and fibrosis, whereas Lira significantly ameliorated these adverse effects in HFD rats. Metabolomic results indicated that Lira could directly reduce renal lipid and energy metabolic disorders in HDF rats. Furthermore, we revealed that Lira inhibited renal lipids deposition by coordinating lipogenic and lipolytic signals, and restored renal mitochondria function via Sirt1/AMPK/PGC1α pathways in HDF rats. Our findings suggested that GLP-1 receptor agonist is a promising therapy for obesity-associated CKD.

## Acknowledgments

This work was supported by grants from National Natural Science Foundation of China (31200754, 81571808) and China Postdoctoral Science Foundation (2012M511931).

## Author Contributions

J.P.L. and Y.R.L. contributed to experimental design. C.S.W., L.L., S.Y.L. and G.N.L. contributed to animal experiments. C.S.W., Y.N.C., J.Q.C. and J.P. L. contributed to sample analysis and result interpretation. C.S.W., J.P.L. and Y.R.L. wrote the paper.

